# Predicting dynamic cellular protein-RNA interactions using deep learning and *in vivo* RNA structure

**DOI:** 10.1101/2020.05.05.078774

**Authors:** Lei Sun, Kui Xu, Wenze Huang, Yucheng T. Yang, Lei Tang, Tuanlin Xiong, Qiangfeng Cliff Zhang

## Abstract

Interactions with RNA-binding proteins (RBPs) are crucial for RNA regulation and function. While both RNA sequence and structure are critical determinants, RNA structure is dependent on cellular environment and especially important in regulating dynamic RBP bindings across various conditions. However, how distinct it contributes to RBP binding *in vivo* remains poorly understood. To address this issue, we obtained transcriptome-wide RNA secondary structure profiles in multiple cell-types, and established a deep neural network, PrismNet, that uses in *vivo* RNA structures to accurately predict cellular protein-RNA interactions. With a deep learning “attention” strategy, PrismNet discovers the exact binding nucleotides and their mutational effect. The predicted binding sites are highly conserved and enriched for rare, deleterious genetic variants. Remarkably, dynamic RBP binding sites are enriched for structure-changing variants (riboSNitches), which are often associated with disease, reflecting dysregulated RBP bindings. Our resource enables the analysis of cell-type-specific RNA regulation, with applications in human disease.

**Highlights:** 1, A big data resource of transcriptome-wide RNA secondary structure profiles in multiple cell types

2, PrismNet, a deep neural network, accurately models the sequence and structural combined patterns of protein-RNA interactions *in vivo*

3, RNA structural information *in vivo* is critical for the accurate prediction of dynamic RBP binding in various cellular conditions

4, PrismNet can dissect and predict how mutations affect RBP binding via RNA sequence or structure changes

5, RNA structure-changing RiboSNitches are enriched in dynamic RBP binding sites and often associated with disease, likely disrupting RBP-based regulation

## Introduction

RNA binding proteins (RBPs) play essential roles in regulating the transcription, metabolism, and translation of cellular RNAs (Baltz et al., 2012; Castello et al., 2012; Gerstberger et al., 2014; Licatalosi and Darnell, 2010). Determining RBP binding profiles in different conditions and elucidating their detailed regulatory mechanisms are critical for understanding their functions. However, given the sheer number of RBPs that accounts for close to 10% of the human proteome (Brannan et al., 2016; Hentze et al., 2018), establishing links between RBPs and their targets has been an enormous challenge. To address this question, many high-throughput technologies have been developed to profile and predict RBP binding. Assays such as systematic evolution of ligands by exponential selection (SELEX), RNAcompete, and RNA Bind-n-Seq can characterize the sequence preferences of RBPs *in vitro* (Ellington and Szostak, 1990; Lambert et al., 2014; Ray et al., 2009), and methods like RNA immunoprecipitation (RIP) and UV crosslinking followed by immunoprecipitation (CLIP) and sequencing can identify RBP binding sites *in vivo* (Gilbert and Svejstrup, 2006; Hafner et al., 2010; Licatalosi et al., 2008; Van Nostrand et al., 2016). Nevertheless, despite extensive mapping of RBP binding, the underlying characteristics remains largely unclear, due to the complexities of high non-specific background, interactions with other RBPs and/or cofactors, and existence of alternative RNA secondary structures of the RNA targets (Darnell et al., 2005; Dominguez et al., 2018; Sun et al., 2019).

In addition to the methods of direct measuring, other approaches have been developed to model and predict RBP binding. Traditionally, position-weight-matrices have been used to describe RBP binding determinants and to predict RBP binding targets from RNA sequences (Bailey et al., 2009). Machine learning methods that integrate different types of information also have been developed to more accurately characterizes the binding pattern of RBPs (Li et al., 2010; Maticzka et al., 2014; Orenstein et al., 2016b). More recently, deep learning (LeCun et al., 2015) approaches have been successfully applied to model protein-RNA interactions and predict RBP binding sites (Alipanahi et al., 2015; Ben-Bassat et al., 2018; Gandhi et al., 2018; Ghanbari and Ohler, 2020; Koo et al., 2018; Zhang et al., 2016). For example, DeepBind was developed to learn RBP binding preferences from RNAcompete data using a deep neural network (Alipanahi et al., 2015).

Although these learning methods successfully capture RBP binding preferences of primary sequence, their prediction accuracy under different physiological states is limited because the RNA sequence is not dependent on *in vivo* conditions. Over the years, several methods have been developed to include RNA structural features of RBP targets in their modeling, but these structures were based on computational prediction rather than *in vivo* analysis (Li et al., 2010; Maticzka et al., 2014; Orenstein et al., 2016b; Zhang et al., 2016). Although RNA structure can be predicted from sequence with some accuracy (Eddy, 2014; Seetin and Mathews, 2012), the predictions do not reflect the dynamic regulations by cellular trans-factors and usually show substantial differences to *in vivo* structures (Rouskin et al., 2014; Spitale et al., 2015). Thus, *in vivo* RNA structure data are essential for accurate modeling and predictions of protein-RNA interactions in physiologically relevant contexts.

Here, we bridge this knowledge gap by determining transcriptome-wide RNA secondary structures in multiple cell types. We then integrate this experimentally-derived information in the construction of a deep discriminative neural network (PrismNet, <u>P</u>rotein-<u>R</u>NA Interaction by <u>S</u>tructure-informed <u>M</u>odeling using deep neural <u>NET</u>work) that accurately models and predicts RBP targets *in vivo*. We apply PrismNet to predict how genomic variants in affect RBP binding, especially in the context of human diseases. Specifically, we focused on single nucleotide variants that disrupt RNA structure (riboSNitches) and often associated with human disease, and discovered that riboSNitches are enriched in dynamic, cell-type-specific RBP binding sites.

## Results

### RNA structuromes in different cell types reveal prevalence of structurally variable sites and their association with dynamic RBP binding

RNA structure is flexible, and this feature plays an instrumental role in determining the varying protein-RNA interactions in different cellular conditions (Gerstberger et al., 2014; Jankowsky and Harris, 2015; Lewis et al., 2017). To elucidate the global relationship between RNA structure and RBP binding, we used *in vivo* click selective 2’-hydroxyl acylation and profiling experiment (icSHAPE) technology (Spitale et al., 2015) to generated a comprehensive resource of RNA secondary structures in seven cell types: K562, HepG2, HEK293, HEK 293T, HeLa, H9 and mES cells (Figures 1A, S1A). These cell lines were selected based on their availability of rich RBP binding data by CLIP-seq (Figure S1A), which allowed for integrative structural-and-interaction modeling of RBPs in a matched cellular context.

**Figure 1.**
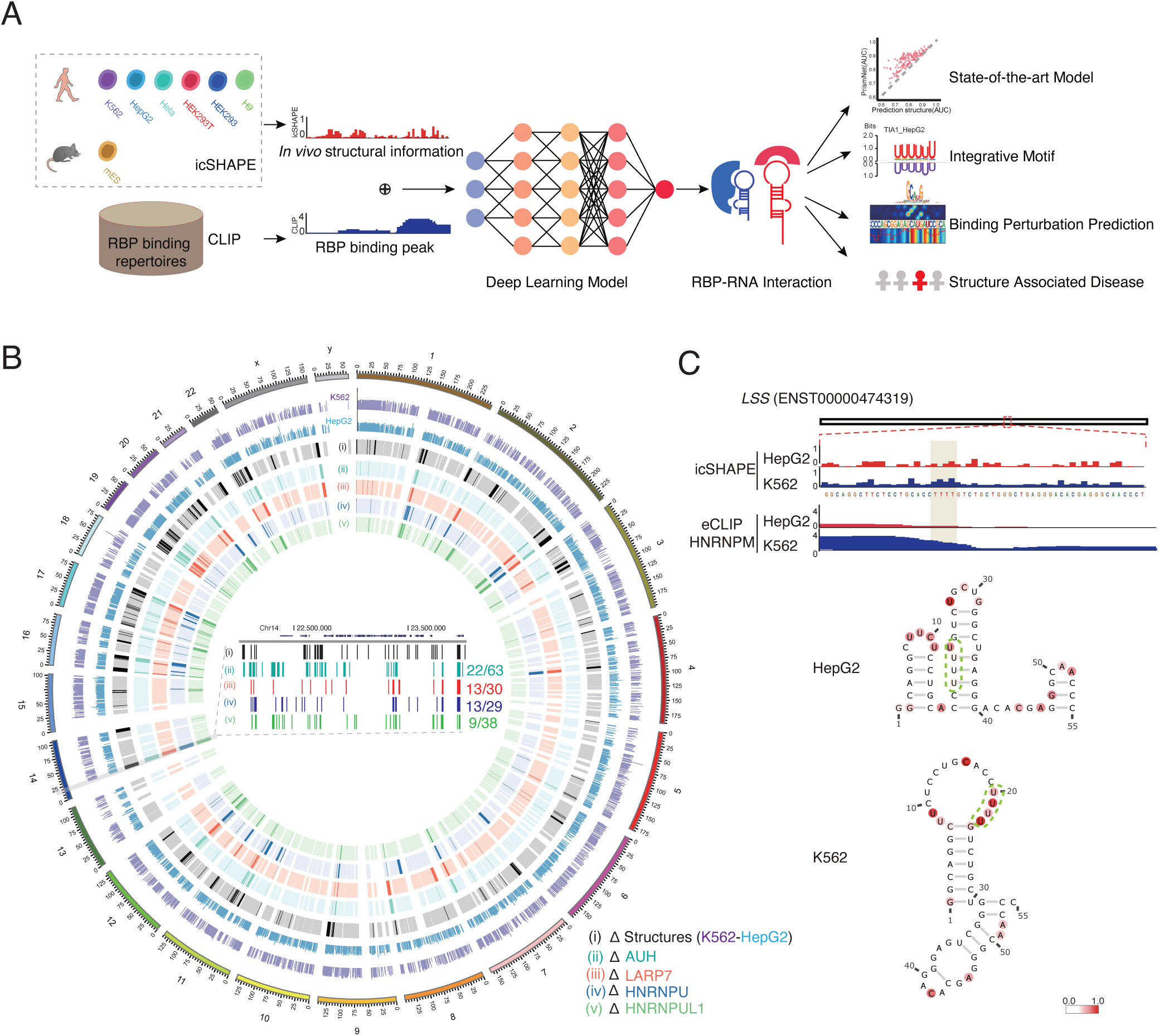
Association between RNA structural variations and dynamic RBP bindings can be used to predict RBP bindings in varying cellular context. **(A)** Integrative modeling and prediction of RBP bindings by PrismNet using *in vivo* RNA structure and RBP binding sites from matched types of cells. PrismNet can be used to dissect and predict the perturbation effects of disease-associated genetic variations on RBP binding. **(B)** Circos plot showing the relationship between RNA structural variations (ΔStructures) and dynamic RBP binding sites (ΔRBP) in HepG2 and K562 cells. The transcripts in the region chr14:22,000,000-24,000,000 is magnified to illustrate that dynamic RBP binding sites is overlapped with RNA structural variations. Numbers show the fraction of overlapped over all RBP binding sites (e.g., 22/63). **(C)** RNA structural and HNRNPM binding profiles in HepG2 and K562 cell lines. Top: icSHAPE scores in the two cell lines for transcripts *LSS*. Middle: Binding site of HNRNPM on *LSS* (eCLIP). Bottom: RNA structural models of the HNRNPM binding sites on *LSS* in the two cell lines. Models are constructed by *RNAshape* with icSHAPE score constraints. Green dashed lines indicate the known HNRNPM poly-U binding motif.

On average, we obtained at least 200 million usable reads for each library replicate after quality control, totaling 4.4 billion reads. We determined RNA secondary structures of the transcripts using icSHAPE-pipe (Li et al., 2019). Our data achieved high coverage of the global transcriptomes (>50,000 transcripts in human; >30,000 transcripts in mouse) as well as high quality (RPKM Pearson correlation coefficient >0.97 between replicates) (Figures S1A-B). For example, our icSHAPE profiling data on 18S rRNA from different human cell lines were highly consistent (Figure S1C) and agreed well with known 18S secondary structures from crystal structures (Figures S1D-E).

Previously, we found that although RNA structure is relatively stable across different subcellular locations, it also contains many structurally variable sites (Sun et al., 2019). These structurally variable sites are hotspots for post-transcriptional regulation, including RBP binding and RNA modification. We found that this is also true when comparing RNA structures across different cell lines, i.e., most of the RNA structures are stable across all cell lines tested, but they also contain a substantial fraction of regions that display structural variability (Figure 1B and S2A-B).

RBP binding could be affected by the diverse cellular environments and thus expected to be dynamic across cell types. We re-analyzed available enhanced CLIP (eCLIP) data and indeed observed very different binding profiles for the same RBPs in different cell lines. For example, on average, only anywhere between ∼15% and ∼60% of the binding sites are shared in K562 and HepG2 cells (Figure S2C). Importantly, we found a high level of association between these dynamic RBP binding sites and the RNA structurally variable sites between the two cell types (Figure 1B, Figure S2D). As an example, HNRNPM is known to bind poly-U sites with single-stranded structure(Van Nostrand et al., 2016), exemplified by the binding sites in the *LSS* and *FAH* transcripts in K562 cells (Figure 1C and S2E). Accordingly, these sites lose HNRNPM binding when switching from the single-stranded to a more double-stranded conformation in HepG2 cells. Overall, these data suggest that RNA structure could play an essential role in determining the dynamic RBP binding in diverse cellular conditions. It is thus critical to incorporate the *in vivo* RNA structural information into platforms that model and predict RBP binding - and their changes - across diverse cellular conditions.

### PrismNet accurately predicts RBP binding *in vivo* using deep learning and RNA structural data

We constructed PrismNet, a deep neural network to accurately model and predict RBP binding, by integrating the *in vivo* RNA secondary structure profiles that we generated with aggregated big data of RBP binding sites. In total, we included 60 RBPs from POSTAR (Hu et al., 2017), 22 RBPs from starBase (Yang et al., 2011), as well as 59 RBPs from ENCODE (Van Nostrand et al., 2016) (Figure S1). Importantly, in contrast to previous methods that have only considered RNA sequences, or relied on computationally predicted RNA structures, PrismNet was constructed to learn protein-RNA interaction determinants from both RNA sequence and *in vivo* structure simultaneously. This approach proved to be vital for capturing the complex interplay between cell type-specific changes in structures and interactions *in vivo*.

For each RBP and an available CLIP experiment, PrismNet trained a model that evaluates every position within the binding sites and calculated a score that reflects the binding contribution of that nucleotide, using the 5,000 most confident peaks from CLIP. As part of the input data, the icSHAPE structure scores of each nucleotide in the same cell type of the CLIP experiment were encoded as a one-dimensional vector, together with the four-dimensional one-hot-encoded sequence (Figure 2A). The PrismNet architecture uses a convolutional layer, a two-dimensional residual block (He et al., 2016) and a one-dimensional residual block connected by max pooling to capture sequence and structural determinants spanning large distances in transcripts. A squeeze-and-excitation (SE) module (Hu et al., 2019) is applied to adaptively recalibrate convolutional channels for learning channel-wise attention (Figure S3A). To mitigate potential overfitting of the PrismNet, we added dropout (Srivastava et al., 2014) layers after every residual block, and other regularizers including weight decay (Hanson and Pratt, 1989) and early stopping in the training stage. Importantly, we also applied SmoothGrad (Wattenberg, 2017) to enable the enhanced saliency maps (Zisserman, 2014) for the visualization and identification of high attention regions (HARs), which are predicted to be the exact locations of RBP binding nucleotides (Figure 2A).

**Figure 2.**
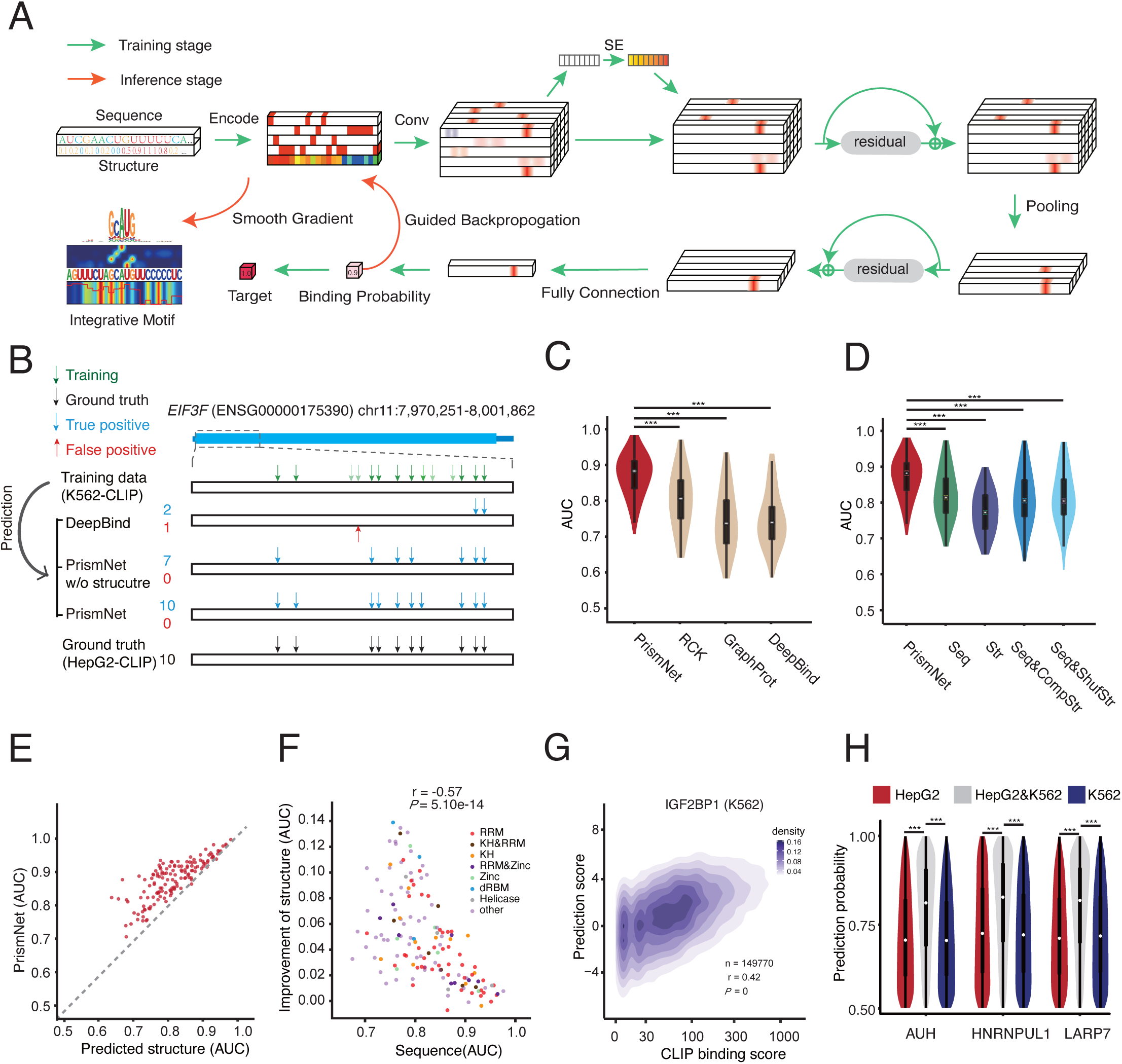
PrismNet more accurately predicts RBP binding in cellular conditions and *in vivo* RNA structure is critical to the prediction improvement. **(A)** Model architecture of PrismNet. The input features include the RNA sequence encoded in the 4-dimensional one-hot encoding and icSHAPE structural score as the fifth-dimension. The neural network consists of multiple convolutional layers, squeeze-and-excitation networks and residual blocks to capture the joint sequence-and-structural determinants of RBP binding. SE, squeeze-and-excitation. **(B)** Predicted versus observed binding sites of IGF2BP1 on the *EIF3F* transcript. Green/black, observed binding sites in K562/HepG2 cells by eCLIP, used as training and ground truth reference data respectively. Blue/red, true/false positive predictions in HepG2 cells based on the models trained on K562 data. **(C)** Violin plot of the overall AUC scores of PrismNet versus other methods in all human 144 PrismNet models of 99 RBPs. \*\*\**P*<0.001 (two-sided paired t-test). **(D)** Violin plot of the overall AUC scores of PrismNet models using different types of input data in all human 144 PrismNet models of 99 RBPs. \*\*\**P*<0.001 (two-sided paired t-test). **(E)** Scatter plot of AUC scores of PrismNet models using *in vivo* structures versus computationally predicted structures. Each dot represents an RBP. **(F)** Scatter plot of AUC improvements of PrismNet versus AUC scores of PrismNet models using only sequence information. Each dot represents an RBP. **(G)** Density map of binding scores of IGF2BP1 predicted using PrismNet versus the observed binding scores from eCLIP experimetns in K562 cells. **(H)** Violin plot of PrismNet-predicted binding probabilities at the binding sites in K562 cells only, HepG2 cells only, or both. \*\*\**P*<0.001 (unpaired t-test).

To demonstrate that PrismNet can accurately model the sequence and structural basis of RBP binding, we performed a proof-of-principle analysis with the RBP IGF2BP1, which plays important roles in regulating RNA stability and localized translation (Huang et al., 2018). Recently, eCLIP uncovered 46,226 IFG2BP1 binding sites in K562 cells (Van Nostrand et al., 2016). We trained PrismNet with this dataset and our icSHAPE data also in K562 cells. We then applied this model to predict the IGF2BP1 binding sites in HepG2 cells, using their corresponding HepG2 icSHAPE data. The predictions were then compared to the HepG2 eCLIP results from the same group, as ground truth. According to eCLIP, the EIF3F transcript contains 14 IGF2BP1 binding sites in K562 cells, and 10 binding sites in HepG2 cells. We found that PrismNet correctly predicted all 10 binding sites within the EIF3F transcript in HepG2 cells with no false positives, by using the model trained in K562 cells. In contrast, DeepBind (Alipanahi et al., 2015), a deep learning model taking sequence only as input, correctly predicted only 2 of the 10 sites (Figure 2B). Interestingly, if we did not include RNA secondary structure data in the training of PrismNet, the sequence-only version correctly predicted only 7 of the 10 sites, indicating that RNA structure information is important for accurate binding site prediction by PrismNet.

We then systematically evaluated the prediction performance of PrismNet by using transcriptome-wide binding sites of all 99 RBPs, and comparing with other state-of-the-art computational methods, including RCK (Orenstein et al., 2016b), GraphProt (Maticzka et al., 2014) and DeepBind (Alipanahi et al., 2015). As mentioned above, DeepBind is the first deep learning model to predict RBP binding sites utilizing only RNA sequences; GraphProt models RBP binding sites based on sequence and predicted structure with graph-kernel features; RCK infers protein-RNA binding preferences using a *k*-mer based model with RNA sequences and predicted structure. For every CLIP-seq dataset of an RBP, we split the binding sites into a training and a test set and use the same sets to benchmark all prediction methods. We observed that, overall, PrismNet achieved the highest prediction performance in terms of Area Under the receiver operating characteristic Curve (AUC) (Figure 2C). For most RBPs, PrismNet constantly showed superior prediction performance (Figures S3B-C). Some RBPs show dramatic performance improvement by PrismNet (e.g., SLTM: PrismNet = 0.82 versus GraphPort = 0.61; XRN2: PrismNet = 0.87 versus GraphPort = 0.62; DGCR8: PrismNet = 0.89 versus GraphPort = 0.63).

### RNA secondary structure *in vivo* is critical input for accurate prediction of RBP binding

To dissect how RNA sequence and structural information contribute to the accurate predictions of PrismNet, we trained the PrismNet model using different combinations of input data: (i) sequence and experimentally-measured structure, (ii) sequence only, experimentally-measured structure only, (iv) sequence and structure predicted by RNAshape (Steffen et al., 2006), and (v) sequence and randomly generated structure. As expected, the model with sequence and experimentally-measured structure (i) as input significantly outperformed other models (Figures 2D and S3D). Notably, PrismNet achieved better performance on almost all RBPs over predictions based on sequence and computationally predicted structures (iv) (Figure 2E). It is also interesting to note that PrismNet could fairly accurately predict protein-RNA binding sites using RNA secondary structure data only (iii), although the prediction accuracy was inferior to that only using sequence data only (ii).

Unexpectedly, we observed comparable prediction performance for the three models that use sequence data only (ii), or sequence data with predicted structures and sequence data with randomly generated structures (v) (Figures 2D and S3D). A recent study showed that predicted structures did not improve the prediction performance of protein-RNA binding if the deep learning models were appropriately designed (Koo et al., 2018). This finding is consistent with our results: a good deep neural network can implicitly retrieve and use sequence-embedded RNA structure information, just like independent predictions. Fledging separately predicted RNA structures thus cannot further improve the prediction (Figure S3E). However, it was surprising that training PrismNet with randomly generated RNA structures did not lead to a visible deterioration of prediction performance, implicating that PrismNet is robust to the noise in the input RNA structural data (Figure 2D).

Interestingly, the prediction improvement brought by experimentally-measured *in vivo* structural data was more significant for those RBPs with lower performance in the sequence-only model (Figure 2F). The level of improvement by the provided RNA structural information was associated with the type of RNA binding domain in the RBPs (Figure 2F and S3F). For instance, RBPs containing single-stranded RNA (ssRNA)-binding domains, such as the RNA recognition motif (RRM) and the K homology (KH) domains, were more dependent on RNA sequence for target recognition, and the prediction improvement from RNA structure was less when comparing to other domains. On the other hand, RBPs containing a double-stranded RNA (dsRNA)-binding motif (dsRBM, *e*.*g*., SND1 and DGCR8) and helicase domains (*e*.*g*., DDX42 and DDX3X) were the least accurate in RNA sequence-only predictions, and the improvement from RNA structural information was the most dramatic. These findings confirm that, in general, RNA structure influences the binding of dsRNA-binding domains more than that of ssRNA-binding domains.

Finally, the predicted binding probability from PrismNet correlated well with the binding affinity determined from CLIP experiments, as shown for different RBPs in K562 cells (Spearman correlation coefficient = 0.46, *P*=2.2e-16) (Figures 2G and S3G-H). Although we only used “1/0” labels to denote the binding and unbinding events in the training dataset, PrismNet apparently learned a quantitative model for RBP binding from the big data of sequence, structure, and protein-RNA interaction. Unexpectedly, cell type-specific binding sites (Figure S2C) generally had lower predicted binding scores compared to common binding sites (Figure 2H, *p* = 0 for unpaired *t*-test). PrismNet is thus able to predict dynamic RBP bindings with intermediate affinity, which likely play important regulatory roles in response to changes in cellular environments.

### RBP binding sites predicted by PrismNet correlates with post-transcriptional regulation

Given that PrismNet could accurately and quantitatively predict RBP binding, we next asked whether the predicted binding sites and binding affinity correlate with RBP-mediated post-transcriptional regulation.

Many of the surveyed RBPs are splicing factors. To investigate the potential concordance between predicted binding sites and alternative splicing, we examined SRSF1, an important splicing factor that functions in both constitutive and alternative pre-mRNA splicing (Pandit et al., 2013). There are over 142,507 SRSF1 binding sites in HepG2 cells, detected by eCLIP (Van Nostrand et al., 2016). We trained PrismNet with this dataset and the HepG2 RNA structural data. PrismNet was then able to predict about 193,584 and 182,886 binding sites in HEK293T and K562 cells, respectively. Of these binding sites, only 133,865 (∼70%) were shared in both HEK293T and K562 cell lines, suggesting a cell-type-specific binding pattern for SRSF1 (Figure S4A).

To determine if dynamic SRSF1 binding sites correlate with alternative splicing, we examined the splicing levels of 12 exons that contain cell-type-specific binding sites in their 5’ splice sites. Indeed, we observed a positive correlation between the differential affinity of the binding sites and differential inclusion levels of these exons in K562 vs HEK293T cells (Spearman correlation coefficient = 0.59, *P*=0.04; Figure 3A, top), suggesting that SRSF1 binding contributes to exon inclusion (Anczukow et al., 2015). We further used RNA-seq data to test all the exons with cell-type-specific SRSF1 binding sites in their 5’ splice sites, and found that the differential affinity of the binding sites was modestly correlated with the differential splicing scores of the exons (Spearman correlation coefficient = 0.183, *P*<4.895e-6; Figure 3A, bottom).

**Figure 3.**
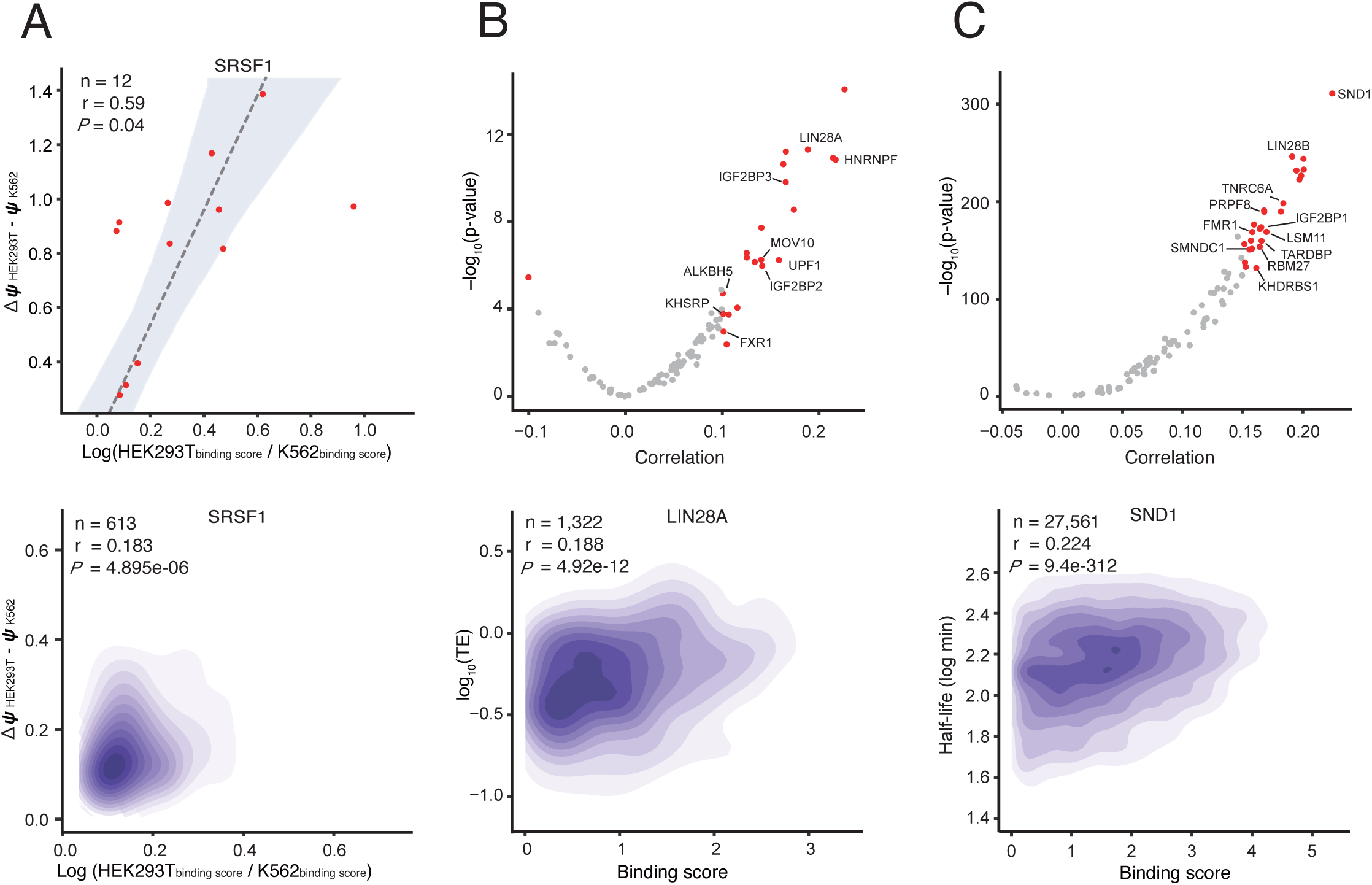
The quantitative RBP binding of PrismNet prediction correlates with the regulations of splicing, translation and degradation. **(A)** Correlation of PrismNet-predicted differential SRSF1 binding and alternative splicing between HEK293T and K562 cells. Top: on experimentally derived alternative splicing scores of 12 exons. Bottom: on transcriptome-wide RNA-seq data. **(B)** Correlation between PrismNet-predicted RBP binding and translational efficiency in HepG2 cells. Top: Distribution of correlation coefficients of all RBPs with significance scores. Marked RBPs are known translation regulators. Bottom: Density plot of the predicted LIN28A binding scores versus translation efficiencies for the target transcripts. **(C)** Correlation between PrismNet-predicted RBP binding and RNA degradation in HEK293 cells. Top: Distribution of correlation coefficients of all RBPs with significance scores. Marked RBPs are known RNA degradation regulators. Bottom: Density plot of the predicted SND1binding scores versus half-lives of the target transcripts.

In addition to alternative splicing, RBPs are also essential regulators of translation and degradation (Gerstberger et al., 2014; Hasan et al., 2014). We explored potential associations between the predicted binding sites and translation efficiency as well as RNA half-life for all surveyed proteins. Our analysis recovered many RBPs known to regulate translation efficiency, including the well-studied IGF2BP proteins (Huang et al., 2018), FXR1 (Vasudevan and Steitz, 2007) and LIN28A (Jin et al., 2011) (Figure 3B). Focusing on LIN28A, a well-known RBP that promotes translation (Jin et al., 2011), we found a positive correlation between the binding score predicted by PrismNet and the translation efficiency of the target transcripts (Figure 3B). Moreover, we also confirmed many proteins that stabilize RNA, such as LIN28B (Hafner et al., 2013), FMR1 (Zhang et al., 2018) and SND1 (Paukku et al., 2008). We looked into transcripts with predicted binding sites of SND1, a protein that is known to regulate RNA half-life (Paukku et al., 2008), and found that they showed increased stability (Figures 3C).

This analysis also identified many putative novel regulators of RNA translation and half-life. For example, we found that ALKBH5 (an m^6^A eraser) may promote RNA translation, consistent with that the elimination of excessively deposited m^6^A could reduce translation efficiency (Slobodin et al., 2017) (Figure S4B). SMNDC1, an alternative splicing regulator, may promote RNA stability, consistent with a previous finding showing that alternative splicing could be coupled with nonsense-mediated mRNA decay (Saltzman et al., 2008) (Figure S4C). Overall, these data indicate that PrismNet, especially its quantitative prediction of the binding affinity, can be used to discover post-transcriptional regulators and targets.

### Verification of the RNA structure code of RBP binding preference

To characterize the binding preferences of RBPs, we developed a computational framework to capture the sequence and structural signatures of each binding site using the saliency map, a technology that has been successfully used in computer vision for visual attention retrieval (Wattenberg, 2017). The important regions in each RBP binding site manifested as high attention regions (HARs) in the saliency map, with the quantification of the contribution of every position to the binding. As a proof-of-concept, we visualized the HARs of a splicing regulator, RBFOX2. The highest attention regions (red, Figure S5A) indicate that a change at these positions - in sequence or structure - would most dramatically change RBFOX2 binding probability. Our maps clearly revealed that HARs were enriched for “GCAUG” within single-stranded structures, consistent with previous studies of RBFOX2 (Jin et al., 2003).

The HAR is a theoretical model that quantifies the sequence and structural contributions of a nucleotide to RBP binding. It thus is important to separately assess their roles by experiments. Previously, experimental validations of a binding motif mainly consider how a sequence mutation affect RBP binding (Alipanahi et al., 2015; Grønning et al., 2019), so here we focused on how a structural change will also influence RBP binding on top of sequence.

We first used different melting-and-folding treatments to perturb RNA structure without altering sequence. Briefly, for a given RBP, we selected PrismNet-predicted RNA binding sites that form hairpin structures (both *in vivo* and *in vitro*), with the HAR residing on the stem. We heat-denatured the RNA fragment of the binding site, and then either slow-cooled to allow refolding into the hairpin structure or fast-cooled to retain single-stranded conformation (Li et al., 2008; Liu et al., 2015b). For example, PrismNet predicted a double-stranded binding site for SND1 in the transcript encoding eukaryotic translation initiation factor 1 (EIF1) in human K562 cells (Figure 4A). We found that SND1 showed stronger affinity to the slow-cooled vs. fast-cooled RNA fragment, consistent with the higher affinity of SND1 for the double-stranded conformation (Figure 4A). Using this approach, we also validated the structural preferences of TIA1 for single-stranded conformation (Figure 4B). These data validated that PrismNet accurately predicted the structural preference of SND1 and TIA1. Given that the structural context of the HARs predicted by PrismNet generally depend on the conditions, it again confirms the essential importance to integrate the *in vivo* structural information in RBP binding site prediction.

**Figure 4.**
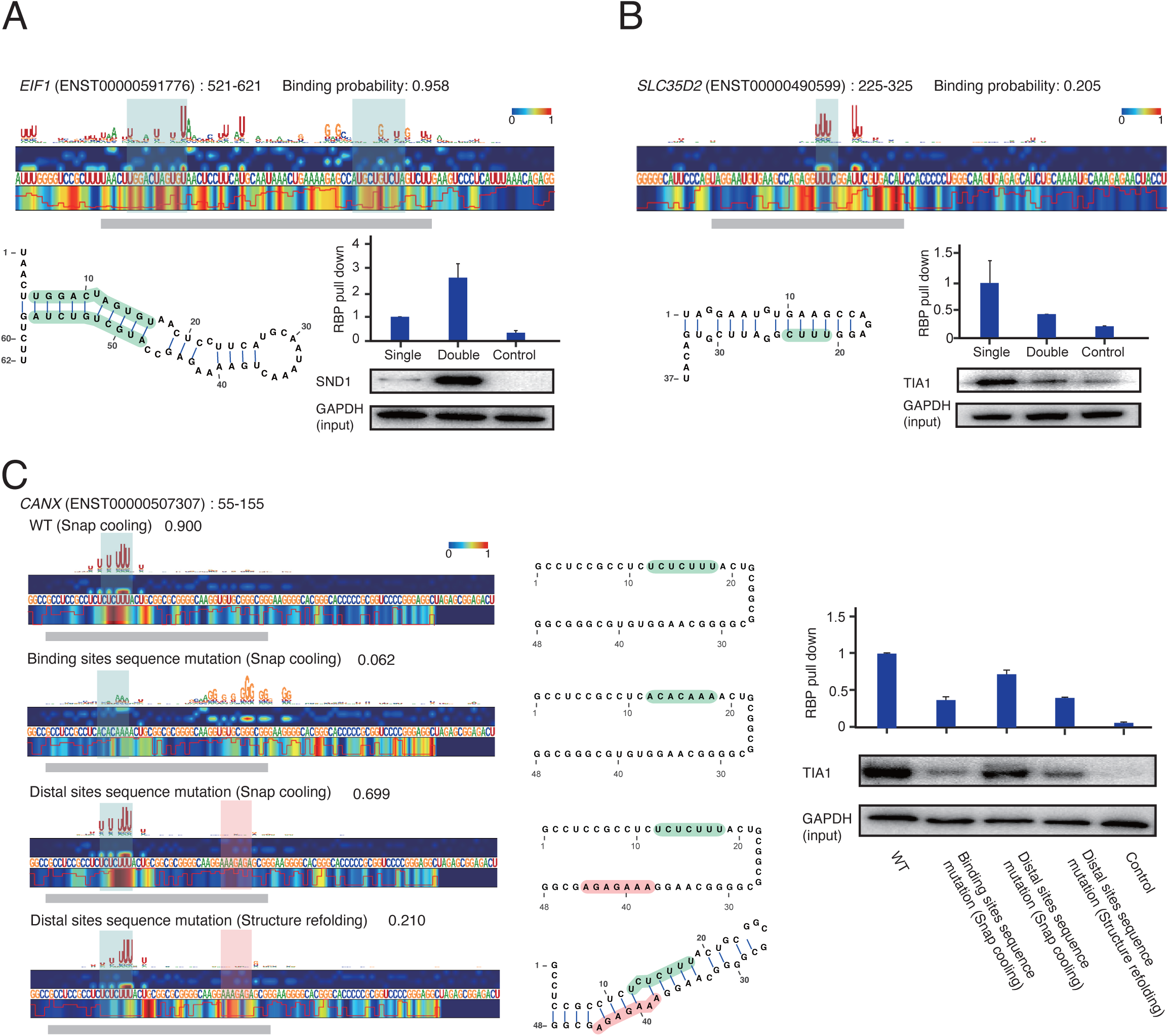
Validation of PrismNet-predicted effects of RNA sequence mutations and structure changes on RBP binding. **(A)** Experimental validation of SND1 binding on a site from the *EIF1* transcript (*EIF1*:521-621). Top: The saliency map showing the site with double-stranded conformation, predicted to be bound by SND1. Grey bar, region of synthesized RNA fragment. Green background, duplex region. Bottom left: Predicted *in vitro* secondary structural model. Bottom right: RNA pull-down for the synthesized RNA fragment in different structural conformation (n=3 replicates). **(B)** Experimental validation of TIA1 binding on a site from the *SLC35D2* transcript (*SLC35D2*:225-325). Top: The saliency map showing the site with double-stranded conformation, predicted *not* to be bound by TIA1. Grey bar, region of synthesized RNA fragment. Green background, TIA1-bindng sequence motif. Bottom left: Predicted *in vitro* secondary structural model. Bottom right: RNA pull-down for the synthesized RNA fragment in different structural conformation (n=3 replicates). **(C)** Experimental validation of TIA1 binding on a site from the *CANX* transcript (*CANX*:55-155). Left: The saliency maps showing the site of the wildtype (WT) and sequence mutants in different structural conformation: (i) WT in single-stranded conformation; (ii) HAR sequence mutation, single-stranded conformation, (iii) distal site sequence mutation, single-stranded conformation, (iv) distal site sequence mutation, double-stranded conformation. Grey, region of synthesized RNA fragments. Green background, TIA1-bindng sequence motif. Pink background, a distal site. Middle: Predicted *in vitro* secondary structural models. Right: RNA pull-down for the synthesized RNA fragment in different structural conformations (n=3 replicates).

As an orthogonal approach to investigate the effect of structural context of HAR, we perturbed the structure by mutating a distal sequence outside of the binding nucleotides. TIA1 is known to bind a poly-U sequence motif (Meyer et al., 2018), and PrismNet discovered that the motif must be in a single-stranded region for high affinity binding. We synthesized a host sequence fragment in the *CANX* mRNA containing a TIA1 HAR. Mutating the poly-U motif disrupted TIA1 binding, even when the whole sequence was kept single-stranded by snap-cooling, confirming the importance of the poly-U sequence for TIA1 binding. In contrast, a mutation at a distal site resulted in diminished TIA1 binding only under conditions that promote base-pairing with the poly-U sequence, further confirming that TIA1 preferentially binds single-stranded poly-U sequences (Figure 4C). Overall, these results verify that PrismNet has learned the regulatory code encoded in RNA secondary structure, thus allowing one to predict how structural alteration affects RBP binding.

### Integrative motifs from PrismNet reveal mechanisms of RBP-RNA recognition

We aggregated and aligned all high attention regions (HARs) to obtain the binding patterns for each RBP, defined as sequence-and-structure integrative motifs (“integrative motifs”). We calculated the integrative motifs for all the RBPs we had surveyed, and used a combined logo to denote a motif by adding a structural component to the nucleotide code (“P” for paired, “U” for unpaired) (Hofacker et al., 1994). The sequence component of the integrative motifs that we discovered were highly consistent with the known sequence preference of these RBPs as documented in the ATtRACT database (Giudice et al., 2016) (e.g. GGA motif for SRSF1, poly-U motif for U2AF2), and with those derived from sequencing experiments by using *k*-mers enrichment (Van Nostrand et al., 2018), or a recent method mCross (Feng et al., 2019) (Figures 5A and S5B-C).

**Figure 5.**
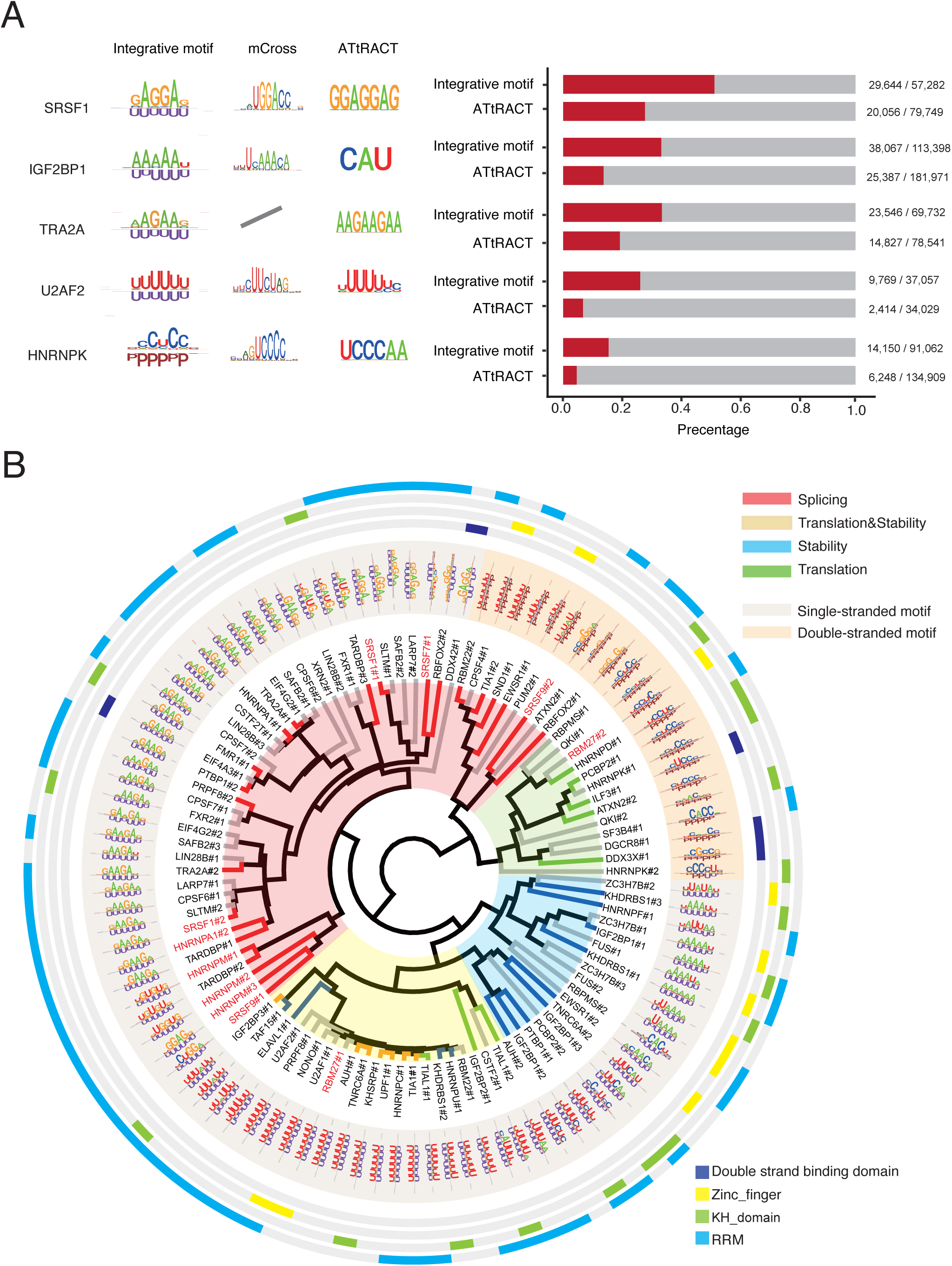
Integrative motifs derived from PrismNet are more specific in capturing transcriptome-wide binding site and clusted with similar post-transcriptional regulatory functions. **(A)** Motifs of different RBPs. Left: the integrative motifs from PrismNet, sequence motifs from mCross and ATtRACT. The integrative motif consists of a sequence component (up) and a structural component (down), where “U” for unpaired and “P” for paired nucleotides. Right: The true positive and all matched RBP binding sites on the transcriptome by motif scanning using the integrative and the ATtRACT motifs. True positives are determined by comparing with eCLIP experiments. **(B)** Hierarchical clustering of integrative motifs of all human RBPs. Color represents functions in the inner tree, structural preferences in the integrative motif circle, and RBDs in the outer circles. RBPs with red fonts are discussed in the main text.

In addition to sequence preferences, all RBPs that we surveyed exhibited preferences for specific RNA structures (Figure 5A). Most RBPs preferred unpaired regions, consistent with previous studies (Dominguez et al., 2018; Li et al., 2010). Importantly, with the structural constraints, integrative motifs had a much lower false positive rate in discovering true RBP binding sites compared to sequence-only motifs (Figures 5A and S5D). For example, respectively 52% (29644/57282) and 25% (20056/79749) of the sites matched by the integrative motif and the sequence-only motif were covered by the experimental CLIP-seq data.

To systematically visualize and compare the sequence and structural specificities for each RBP, we performed hierarchical clustering of all associated integrative motifs. Interestingly, RBPs involved in the same RNA regulatory pathway were generally clustered together (Figure 5B), which can be partially explained by their similar integrative motifs. For example, RBPs preferring the GA (AG/GAA) motifs in single-stranded RNAs were enriched for regulators of alternative splicing and were clustered together, including SRSF family proteins (e.g. SRSF1, SRSF7 and SRSF9) as well as some hnRNPs (e.g. hnRNPA1, hnRNPM). Many RBPs that bind to U-rich, A-rich or AU-rich motifs in single-stranded RNAs were associated with RNA stability and also clustered together (Figure 5B), consistent with previous studies showing that the AU-rich elements (AREs) in 3’ UTRs target host mRNAs for rapid degradation (Vasudevan and Steitz, 2007).

Some RBPs have multiple binding patterns, i.e., integrative motifs. For example, SRSF9 can bind to GUGGA in single-stranded structures and CCGGGA in double-stranded structures, and RBM27 can bind to polyU sequence in single-stranded structures and UCCUC in double-stranded structures. Generally these proteins contain multiple RBDs with different binding preferences and plays diverse regulatory roles (Dominguez et al., 2018; Ray et al., 2013). Overall, these observations suggest that integrative motifs could capture the determinants of RBP binding more accurately and agree with the domain composition and biological functions of the associated RBPs.

### High attention regions predicted by PrismNet represent evolutionarily conserved, functionally important sites

An integrative motif of an RBP displays certain levels of flexibility in sequence and/or structural contents at each position (Figure 5B). Variations beyond the tolerance level may disrupt RBP binding and consequently lead to dysregulation. Indeed, using both the PhyloP and the PhastCons scores from UCSC (Pollard et al., 2010; Siepel et al., 2005), we found that the HARs predicted by PrismNet are more evolutionarily conserved than the overall transcripts (Figures 6A and S6A).

**Figure 6.**
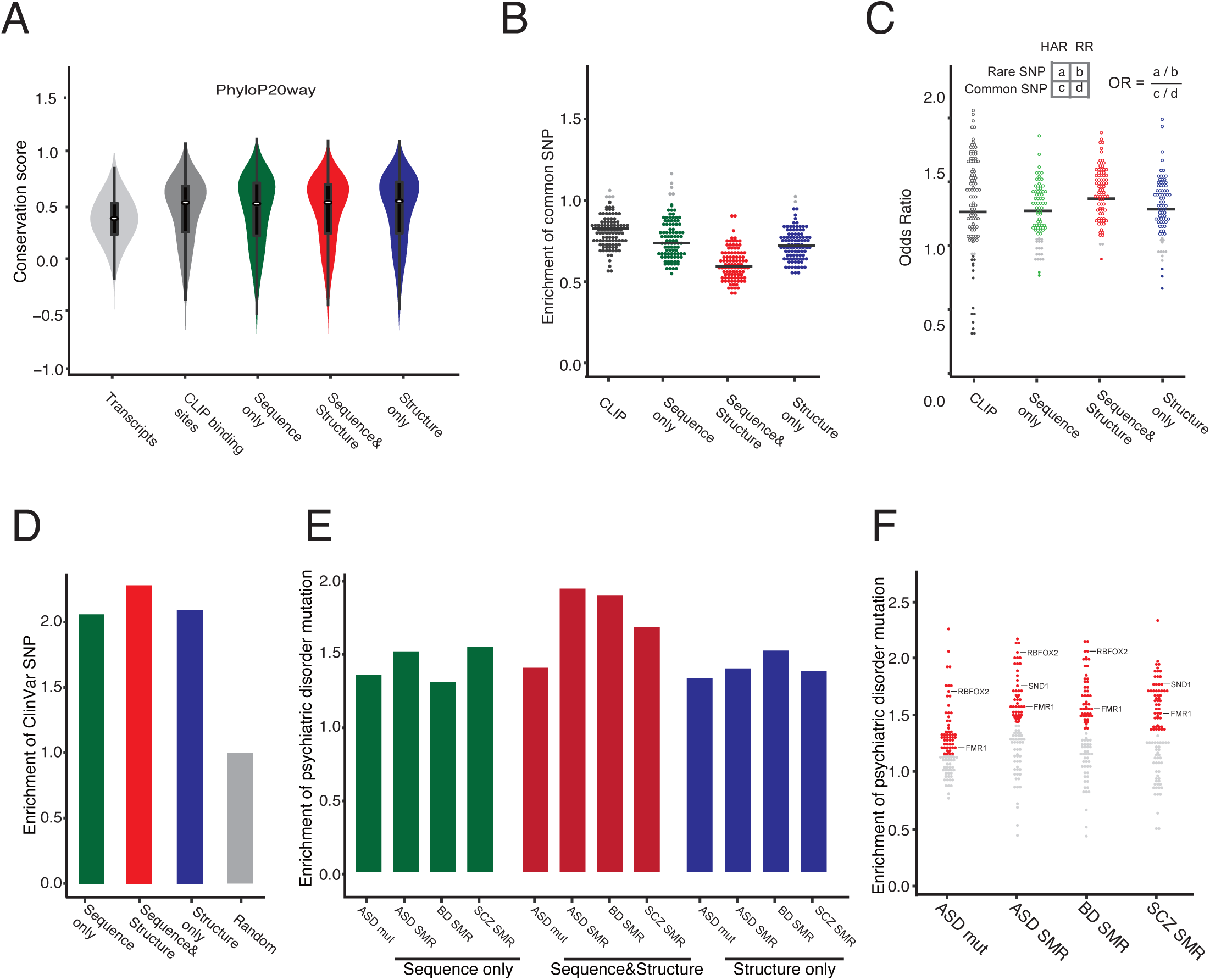
Conservation and enrichment of genomic variants in PrismNet-predicted high attentione regions (HARs) **(A)** Distribution of PhyloP20way conservation scores of HARs, with those of all CLIP binding sites and all transcribed regions as positive and negative controls. **(B)** Enrichment of common SNPs (from dbSNP) in HARs and CLIP binding sites for every RBP comparing to random transcript regions. **(C)** Enrichment of rare SNVs (from dbSNP) relative to common SNVs in HARs and CLIP binding sites for every RBP. **(D)** Enrichment of ClinVar mutation in HARs comparing to random transcript regions. (**E-F**) Enrichment of ASD, BD and SCZ-associated variants in all HARs collectively (**E**) or HARs of every RBP respectively (**F**), comparing to random transcript regions. The RBPs known to associate with psychiatric disorders were highlighted. All HARs and CLIP-seq binding sites are in K562 cells. In **B, C**, and **F**, each dot represent an RBP. RBPs with HARs significantly depleted/enriched of genomic variants (including common and rare SNVs and disease-associated variants) are shown in color (*for P<0.05, permutation or fisher exact test).

SNP density is also an indicator of evolutionary conservation, and has been used to identify constrained regions across the human genome (Sabeti et al., 2007). We analyzed the enrichment of SNPs at HARs versus the whole transcript. First of all, we separated the HARs into three groups according to their attention in primary sequence and/or secondary structure. Briefly, for each binding site, the regions with both high attention sequence and structure components in PrismNet belong to the “sequence & structure” group; the regions with only high attention sequence or structure component are in the “sequence only” or “structure only” group. We then calculated an odds ratio for each group to represent the enrichment of SNPs in HARs over background random positions for every RBP.

We first considered the common SNPs in the dbSNP database (Sherry et al., 2001) and found that all three groups showed a depletion of common SNPs for most RBPs (Figure 6B). Furthermore, while the groups of “sequence only” and “structure only” showed comparable odds ratios, the group of “sequence & structure” were generally more depleted of SNPs (odd ratio: 0.60 (sequence & structure) vs 0.78 (sequence) or 0.75 (structure), *P* < 0.001, two-sides *t*-test). We confirmed these results using the common SNPs from the 1000 Genomes (Altshuler et al., 2015) (Figures S6B).

Next, we investigated the enrichment of rare variants, which, on the contrary, are considered to be deleterious in human populations (Lek et al., 2016). We found that the HARs of most RBPs are enriched for rare SNVs (Figures 6C and S6C), suggesting that the variants within HARs tend to disrupt the functionality of the predicted binding sites. And again, “sequence & structure” group shows higher enrichment than the other two groups. Together, these data suggest that the HARs, particularly with both sequence and structure signatures, are evolutionary conserved, depleted of common SNPs, with harbored variants tending to be deleterious rare mutations.

We thus further explored the relationship of variants in HARs with human disease. Indeed, genetic mutations deposited in the ClinVar database are enriched in HARs, especially in the “sequence & structure” group (Figure 6D). Recently, large-scale neurogenomic studies have identified a large number of eQTL SNPs and *de novo* mutations associated with psychiatric disorders including autism spectrum disorder (ASD), schizophrenia (SCZ), and bipolar disorder (BD) (An et al., 2018; Gandal et al., 2018). In these studies, the effects of the identified variants were mostly explained by their ability to alter chromatin structure and TF binding. Interestingly, we found that these variants were also more enriched within the HARs (Figures 6E). The odds ratios varied for RBPs within each disorder, with some specific RBPs exhibiting stronger enrichment (Figure 6F). For example, RBFOX2, SND1 and FMR1 binding sites were frequently disrupted by ASD-associated variants, consistent with the roles of these proteins as revealed by previous studies (Hagerman et al., 2011; Martin et al., 2007; Weyn-Vanhentenryck et al., 2014). Notably, again, the “sequence & structure” HAR group shows higher levels of enrichment for these psychiatric disorder-associated variants. Thus, analyzing mutations that are within the PrismNet HARs could nominate putative RBP regulators and targets in complex human disorders.

### RNA-structure-disruptive variants (RiboSNitches) are associated with dynamic RBP binding and disease

To further investigate the relationship between mutations in the predicted HARs and human disease, we focused on riboSNitches, a special class of SNPs or SNVs in which different alleles exhibit distinct foci RNA structures (Halvorsen et al., 2010; Wan et al., 2014b) (Figure S7A). RiboSNitches are typically identified by allele-specific structural analysis in the same cells or cells from closely-related individuals (Wan et al., 2014b). Here we developed a novel pipeline to uncover potential riboSNitches by comparing RNA structures at SNP sites with different alleles, and identified thousands of putative riboSNitches for each pair of cell lines (Figure 7A). We intersected this dataset with the riboSNitches identified in a previous study in human lymphoblastoid cell lines (Wan et al., 2014b) and found a significant overlap (Figure S7B), confirming the validity of the dataset.

**Figure 7.**
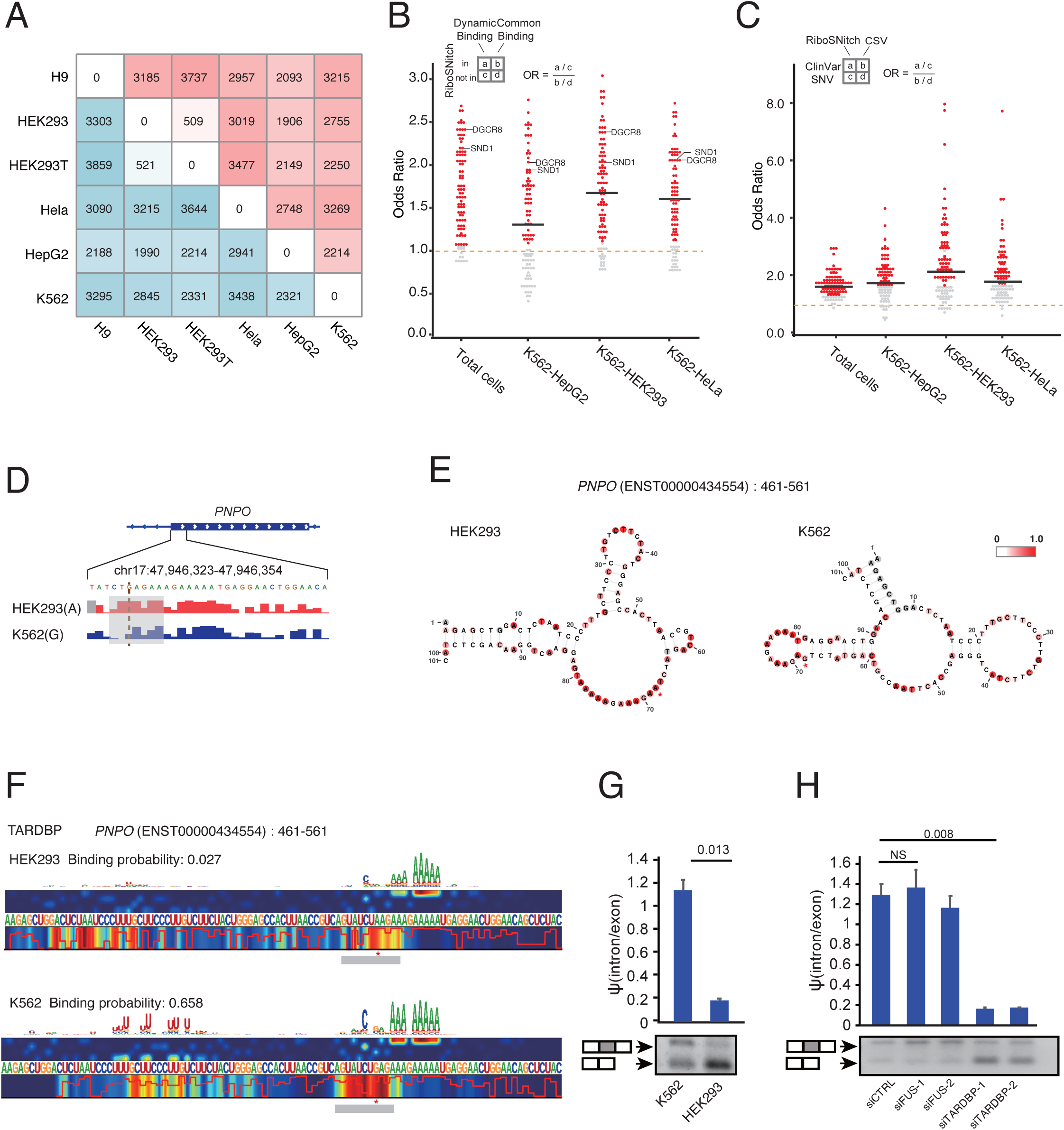
RiboSNitches are associated with RBP dynamic binding sites and human diseases. **(A)** The number of all riboSNitches between different cell lines (bottom-left triangle), and those within dynamic RBP binding sites predicted by PrismNet (up-right triangle). **(B)** Enrichment of riboSNitches in dynamic RBP binding sites comparing to common binding sites predicted by PrismNet. **(C)** Enrichment of riboSNitches relative to CSVs within HARs in human disease (from ClinVar). In **B** and **C**, each dot represents an RBP, with significantly enrichment shown in color (* for P<0.05, fisher exact test). **(D)** Structural profiles of the icSHAPE scores around a riboSNitch in the *PNPO* transcript in HEK293 and K562 cells. **(E)** Structural models of riboSNitch and flanking sequences in the *PNPO* transcript in HEK293 and K562 cells, predicted by *RNAshape* with the corresponding icSHAPE scores as constraints. Red stars indicate the mutation sites. **(F)** Binding probabilities and saliency maps of TARDBP on the *PNPO* transcript in HEK293 and K562 cells. **(G)** Splicing of the *PNPO* transcript in K562 and HEK293 cells (n=3 replicates). **(H)** Splicing of the *PNPO* transcript in WT and TARDBP knockdown K562 cells. FUS knockdowns are included as a negative control.

Notably, these riboSNitches are strongly associated with dynamic RBP binding (Figures 7A-B). For every RBP in every pair of cell lines, we intersected the riboSNitches with the PrismNet-predicted dynamic binding sites. We found that most of these riboSNitches are located in a dynamic binding site of at least one RBP (Figure 7A), and for most RBPs, their dynamic binding sites are enriched with riboSNitches, particularly for those RBPs with double-stranded structural preference (e.g. DGCR8, SND1) (Figure 7B).

Many of the riboSNitches are also disease-associated variants from the ClinVar database (Landrum et al., 2014). Strikingly, most of these disease-associated riboSNitches are located in dynamic RBP binding sites (Figure S7C). Compared to variants that do not affect RNA structure, riboSNitches were more enriched with disease-associated RBP-binding-disruptive mutations, both at the global level and at the level of individual RBPs as well (Figure 7C). Note that to avoid the possibility of protein truncation or frame shift, here we only considered synonymous variants.

To clarify the regulatory relevance of riboSNitches in the context of disease association, we focused on a putative riboSNitch in the pyridoxamine 5’-phosphate oxidase (PNPO) gene for further validation. This riboSNitch corresponds to a synonymous G-to-A mutation (c.552 G>A; p.184 Leu>Leu) that has been associated with epilepsy, a severe neurological disorder (Richards et al., 2015). We found that HEK293 contains both alleles, whereas K562 contains the G allele (Figure S7D). The locus was indeed a riboSNitch, with very different RNA structures in the two cell lines (Figures 7D-E). We then used PrismNet to scan each RBP for its binding probability at this local region and found that TARDBP had a much stronger binding affinity in K562 cells than in HEK293 cells (Figure 7F binding probability: 0.658 in K562 versus 0.028 in HEK293), consistent with the experimental CLIP-seq data (Figure S7E, K562: GSM2423707, GSM2423708; HEK293: DRX012638, DRX012639). TARDBP has been implicated in pre-mRNA splicing (Tollervey et al., 2011) and we noticed that the *PNPO* exon containing this riboSNitch exhibited a dramatically reduced inclusion rate in HEK293 cells (Figure 7G, *P*=0.013, t-test). Interestingly, knockdown of TARDBP in K562 cells resulted in substantially reduced PNPO exon inclusion, in line with the phenotype of disrupted TARDBP binding in HEK293 cells (Figures 7H and S7F). Collectively, these data indicate that riboSNitches, by disrupting the local RNA structure and regulating RBP binding simultaneously, may function to bridge the disease-associated synonymous mutations to RBP-based post-transcriptional regulation.

## Discussion

A substantial body of work has attempted to map RBP binding profiles to gain a more complete and mechanistic understanding of RNA regulation (Baltz et al., 2012; Castello et al., 2012; Gerstberger et al., 2014; Licatalosi and Darnell, 2010). Target recognition by RBPs has been found to be remarkably precise *in vivo* yet usually involves a limited primary RNA sequence space that is rich in low-complexity motifs (Jankowsky and Harris, 2015; Lunde et al., 2007; Singh and Valcarcel, 2005). Binding site predictions with these low-complexity motifs inevitably contain many false positives. Local RNA structure of RBP binding sites is known to contribute critically to the specificity of RBP recognition, however, previous predictive models have not included information on *in vivo* RNA structure due to limited data availability. Here, we resolved these issues by generating a large dataset of RNA structural profiles across diverse mammalian cell lines. Our RNA structurome data shows that RNA structure displays a significant amount of cell line-dependent variability, strongly suggesting its critical importance in the modeling cell line-specific RBP binding. We implemented these insights into a deep neural network, PrismNet, constructed upon the large amount of cell line-specific RBP binding and RNA structurome data *in vivo* from matched cell lines. PrismNet learns protein-RNA interaction models and thus was able to very accurately predict RBP binding in cellular conditions.

The PrismNet model also allows us to dissect the relative contributions of RNA primary sequence and secondary structure. We verified that the change of RNA structure alone could affect RBP binding, once again highlighting its critical role in the dynamic regulation of RBP binding *in vivo*. Applying PrismNet models will accurately predict how RBP bindings vary together with RNA structural changes in new physiological contexts. Our finding also shows the promise of deep learning models for integrative data analysis and providing biological insights, rather than merely serving as black box classifiers.

By extracting features from a big dataset of binding sites with sequence and structural information, PrismNet learns a quantitative model of specificity determinants using the saliency map, in which the contribution of every position can be visualized. More importantly, with the model, PrismNet can predict mutations with severe consequences on RBP binding, and automatically dissect the perturbation effects resulting from the disruption of normal sequence and/or structure patterning. We identified and validated mutations in the PrismNet high attention regions that are strongly deleterious in the human population, and are implicated with potentially important roles in complex human disorders.

Much work remains to be done to understand post-transcriptional regulation in different physiological contexts, particularly in disease-relevant conditions where mutations affect RNA structure and RBP binding. SNPs with allele-specific RNA structure, i.e. riboSNitches, are an emerging topic of great interest with ample opportunities for discovery. Although some candidates were found associated with diverse human disorders and phenotypes (Wan et al., 2014b), we still do not know why such correlations exist and the underlying mechanisms. Here we comprehensively surveyed their association with dynamic RBP binding and also human disease. It is thus a fascinating hypothesis that these correlations could be mediated by the dysregulation of RBP binding. To this end, more RNA secondary structure profiles from disease-related tissue types will be valuable for dissecting the mechanisms of riboSNitches in a specific disease or phenotype context. Another important and further step would be to discover small-molecule drugs that could potentially target RNA in a structure-specific manner (Warner et al., 2018). Together, the enhanced understanding of the regulatory mechanisms that govern the formation of RNA structure and its interaction with other factors may pave the way to the discovery of novel therapeutic candidates.

## Acknowledgments

We thank members of the Zhang labs for discussion. We thank Meiling Piao for experimental advice, Pan Li for computational advice, and Xiaohua Shen, Yafei Yin, Kehkooi Kee and Nan Wang for the gifts of cell lines. This work is supported by the National Natural Science Foundation of China (Grants No. 31671355, 91740204, and 31761163007). L.S. was supported by the Tsinghua-Peking Center for Life Sciences Postdoctoral Fellowship.

## Author Contributions

Q.C.Z. conceived and supervised the project. L.S. performed icSHAPE experiments and other validation experiments. K.X. and W.H designed the deep learning models. W.H., L.S., and Y.T.Y. analyzed all the results. L.T. assisted with experiments, and T.X. assisted with analysis. Q.C.Z., Y.T.Y., and L.S. wrote the manuscript with inputs from all authors.

---

**Supplemental Figure 1.**
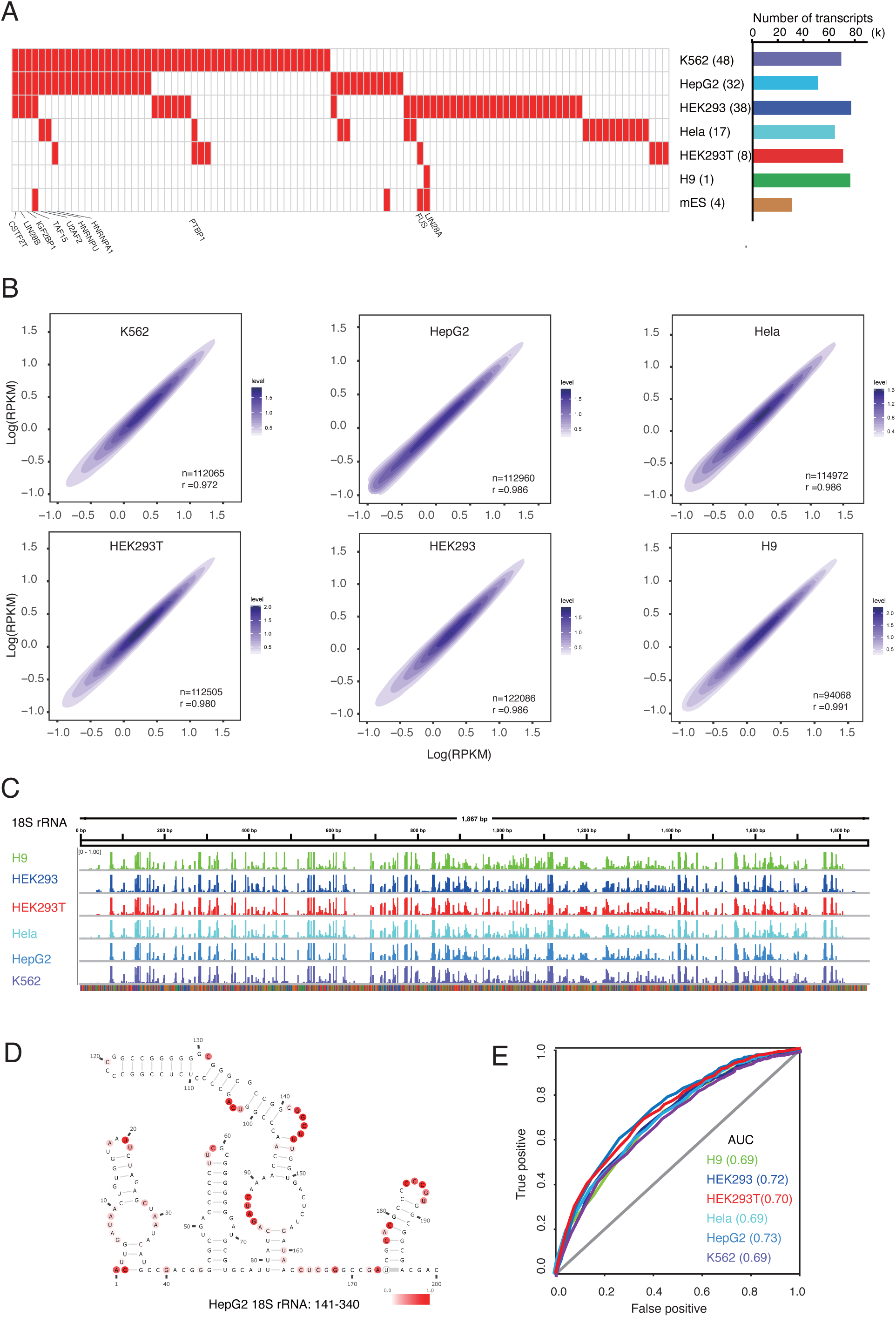
Probing of RNA structuromes by icSHAPE in different cell lines. **(A)** Left: Heatmap of available RBP CLIP datasets in each cell line. RBPs with datasets in at least three cell lines were labeled. Middel: Cell lines with the number of available CLIP datasets. Right: The number of transcripts with secondary structure probed by icSHAPE in each cell line. **(B)** Correlation of RNA expression (RPKM) between replicates of icSHAPE libraries. **(C)** Tracks of icSHAPE scores of the human 18S ribosomal RNA. **(D)** Structural model of 18S rRNA (141-340). Model is ploted with the *ViennaRNA* web service based on data from the *Comparative RNA Web* (*CRW*) Site. Nucleotides are colored with icSHAPE scores from HepG2 cells. **(E)** ROC curve plot of the agreement of icSHAPE scores from different cell lines with the reference 18S rRNA structure from the *Comparative RNA Web* (*CRW*) Site. AUC scores for all ROC curves are included.

**Supplemental Figure 2.**
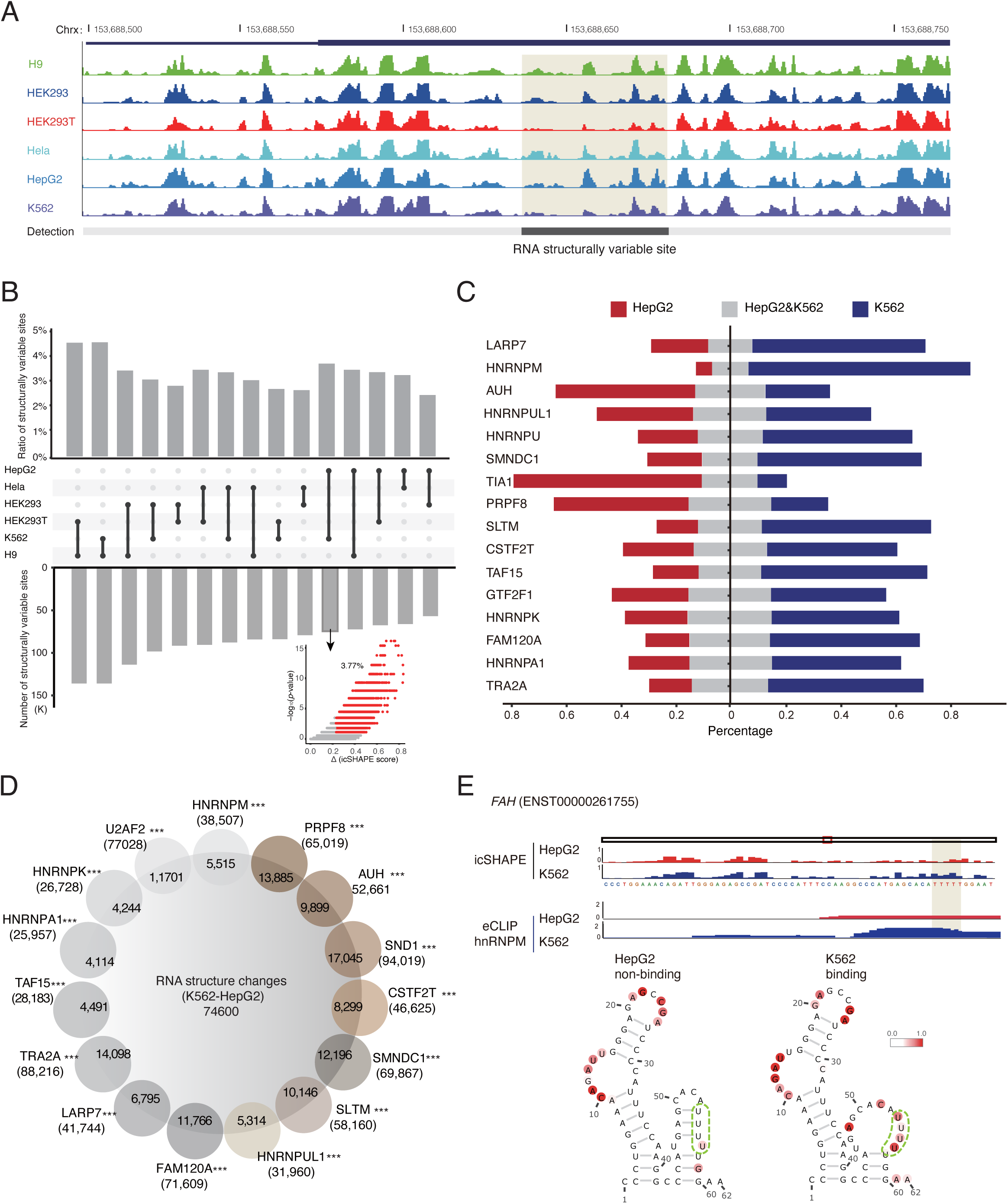
RNA secondary structurally variable sites, dynamic RBP binding profiles, and their associations. **(A)** Tracks of icSHAPE scores at the human genome location chrx:153,688,450-153688850. The light Gray background color labels a structurally variable site. **(B)** Bar plot of the ratio and the number of RNA structurally variable sites between any two human cell lines. Valcano plot shows the statistical significance versus magnitude of RNA structural changes between HepG2 and K562 cells. The red plots are the identified structurally variable sites. **(C)** Stacked bar plots of the percentage of dynamic (cell type-specific) and common RBP binding sites in two cell lines from eCLIP datasets: HepG2-specific (red), K562-specific (blue) and common (grey). **(D)** The overlap between RNA secondary structurally variable sites (big central circle) and dynamic RBP binding sites (small surrdounding circles) in HepG2 and K562 cells.. The total numbers and also the overlapped numbers are shown. *P* values were calculated by fisher exact test. *P < 0.05; **P < 1 ×10^−3^; ***P < 1 × 10^−5^. Note that this is not a Venn diagram in that the overlaps among the RBP binding sites are not considered. **(E)** RNA structural and HNRNPM binding profiles in HepG2 and K562 cell lines. Top: icSHAPE scores in the two cell lines for transcripts *FAH*. Middle: Binding site of HNRNPM on *FAH* (eCLIP). Bottom: RNA structural models of the HNRNPM binding sites on *FAH* in the two cell lines. Models are constructed by *RNAshape* with icSHAPE score constraints. Green dashed lines indicate the HNRNPM binding motif.

**Supplemental Figure 3.**
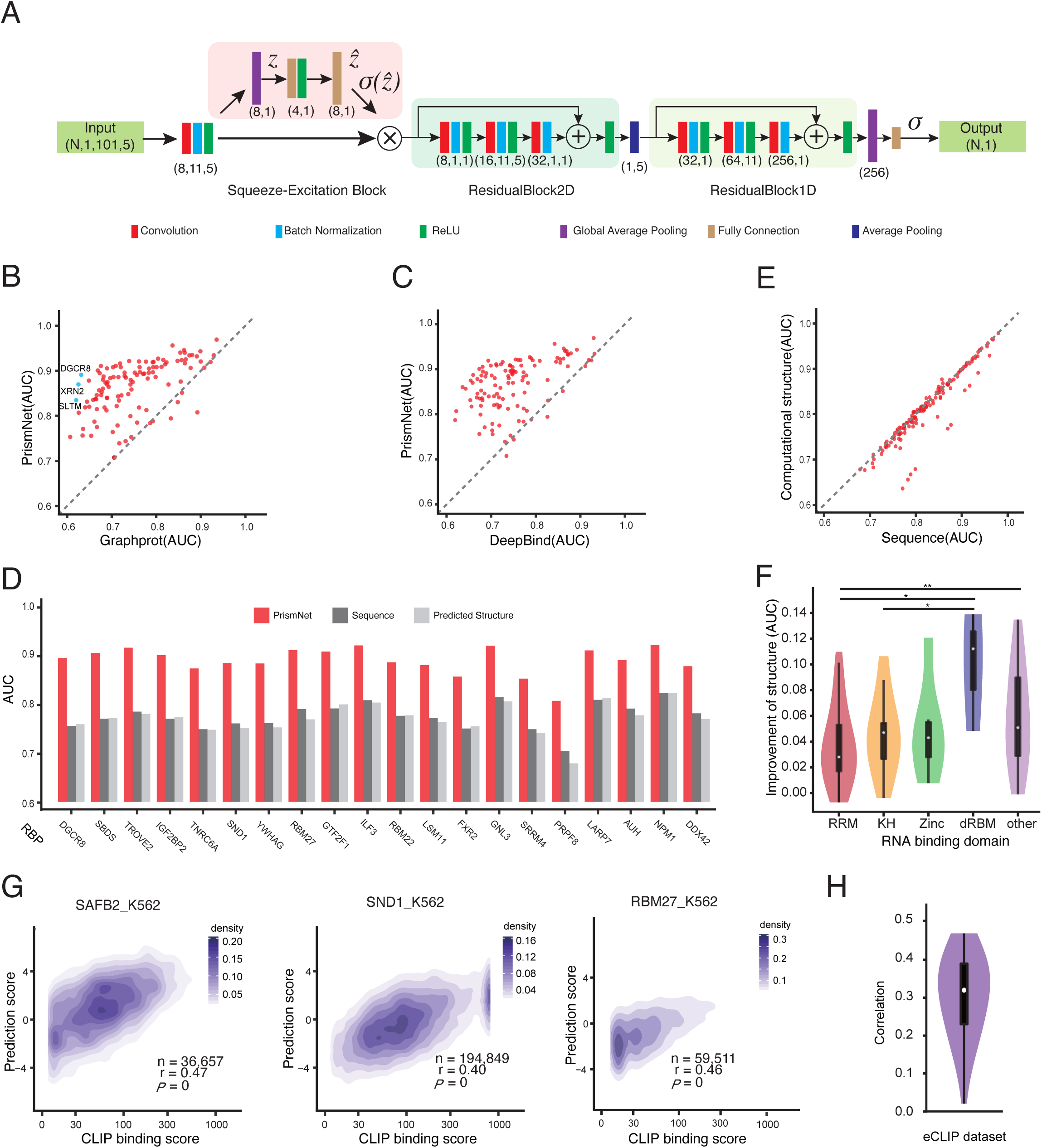
The architechture and prediction performance of PrismNet. (**A**) The architecture of PrismNet with important hyper-parameters. (**B-C**) Scatter plot of AUC scores of the predictions by PrismNet versus GraphPort (**B**) and DeepBind (**C**). Each dot represents an RBP. **(D)** Scatter plot of AUC scores of the predictions by PrismNet with computationally-predicted structure versus with only sequence. Each dot represents an RBP. **(E)** Bar plot of AUC scores of the predictions by PrismNet (red), PrismNet with only sequence (dark grey), and PrismNet with computationally-predicted RNA structure (light grey). **(F)** Violin plot of AUC improvements of PrismNet predictions using *in vivo* structure for RBP groups with different RNA-binding domains. \**P* < 0.05; \*\**P* < 0.01; \*\*\**P* < 1 ×10^−3^ (two-sided unpaired t-test). **(G)** Density plot of the binding probability of SAFB2, SND1, and RBM27 predicted by PrismNet versus the observed binding scores from eCLIP in K562 cells. **(H)** Violin plot of the correlation between PrismNet-predicted binding probability and CLIP binding scores from eCLIP for different proteins.

**Supplemental Figure 4.**
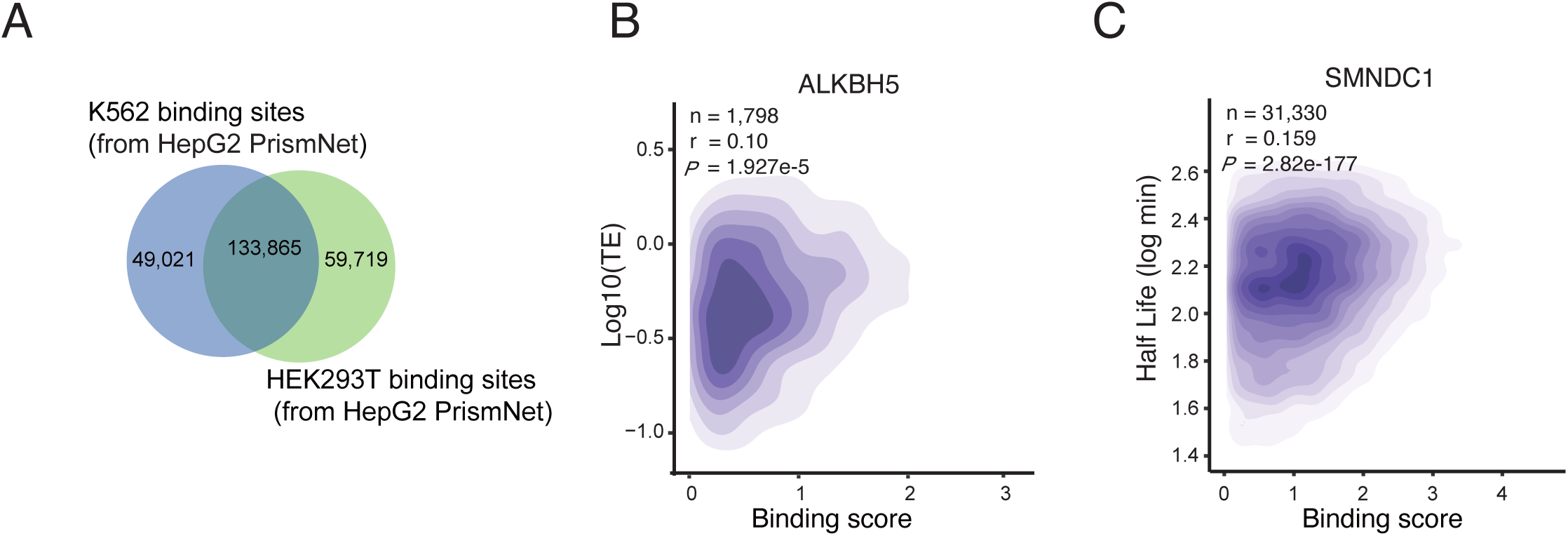
The PrismNet-predicted differential RBP binding, and the correlation of predicted quantitative RBP binding and target translation and degradation. **(A)** Venn diagram of SRSF1 binding sites in HEK293T and K562 cells, predicted by PrismNet model trained on the HepG2 SRSF1 eCLIP dataset. **(B)** Density plot of the PrismNet-predicted ALKBH5 binding scores versus translation efficiency of the target transcripts. **(C)** Density plot of the PrismNet-predicted SMNDC1 binding scores versus half-lives of the target transcripts.

**Supplemental Figure 5.**
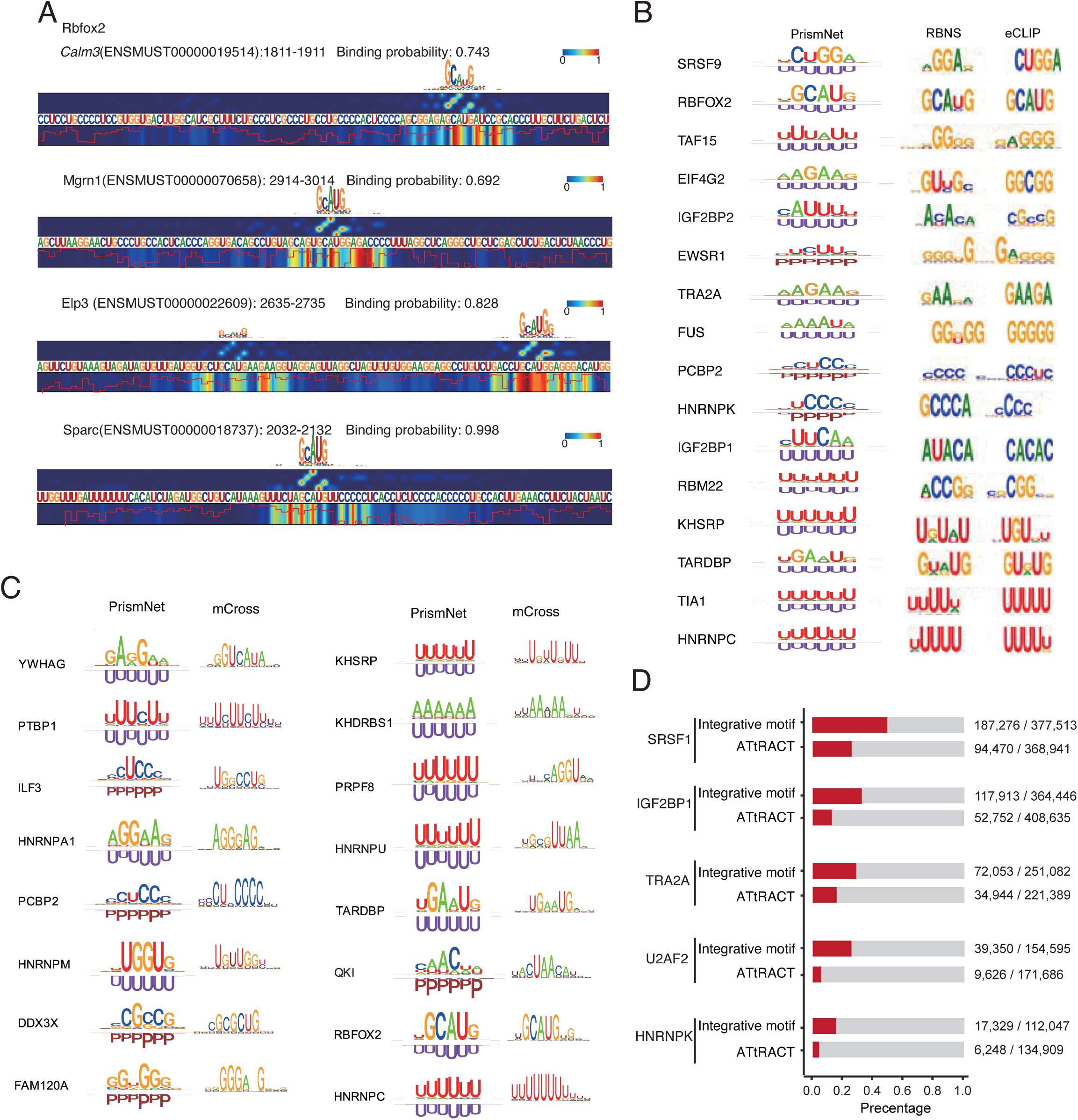
Saliency maps of PrismNet-predicted binding sites and Integrative motifs of different RBPs. **(A)** Saliency maps of Rbfox2 binding sites derived from the mES PrismNet model. The heatmap tracks display the reponse of the model at each nucleotide, with red color showing high attention (upper: sequence reponse, bottom: structure reponse). The sequence logos on the top are shown to help to display high attention sequence component. The red line at bottom represents icSHAPE scores. **(B)** Integrative motifs derived from PrismNet, compared with those provided in eCLIP and RBNS. **(C)** Integrative motifs derived from PrismNet, compared with those obtained by mCross. **(D)** True positives and all matched binding sites on the transcriptome by motif scanning using the integrative and the ATtRACT motifs of different RBPs. True positives are determined by comparing with eCLIP experiments. Related to main Figure 5A. Here we used a loose criterion in motif scanning to match more sites.

**Supplemental Figure 6.**
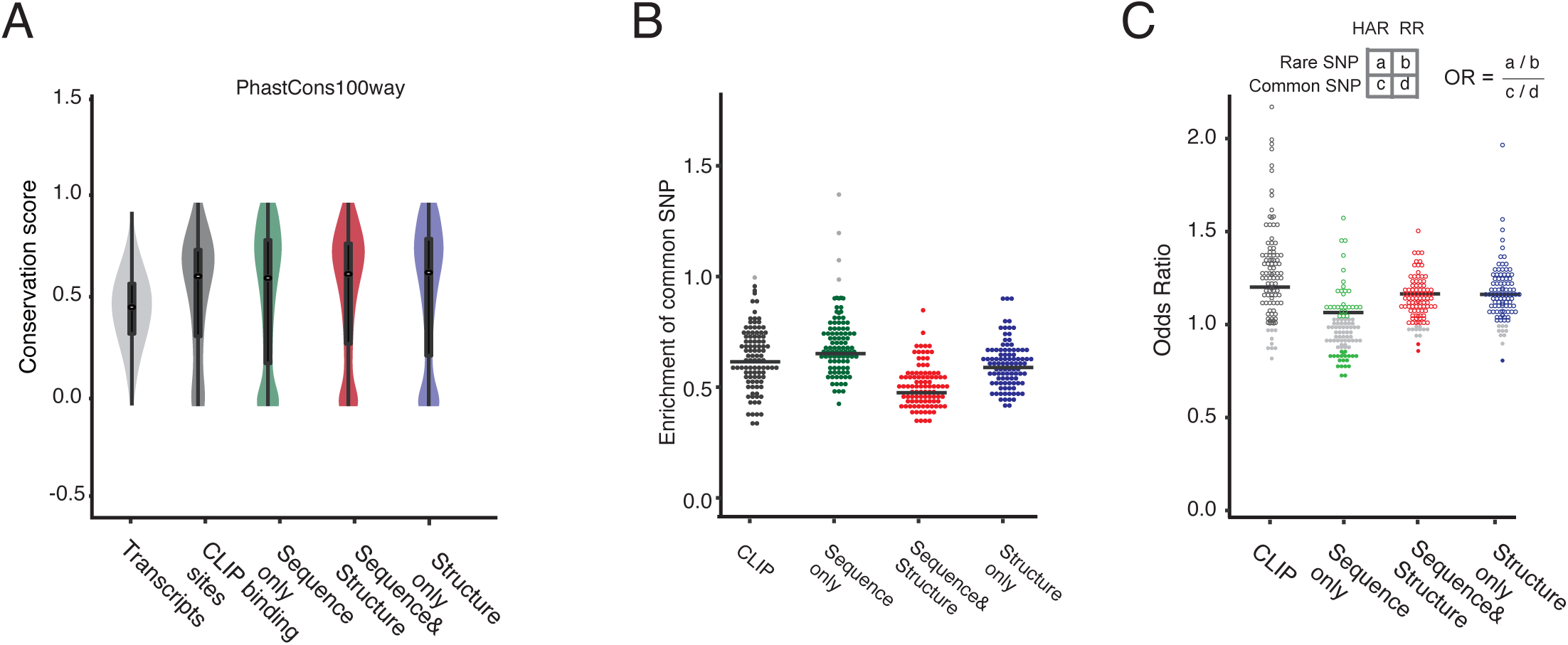
Conservation and enrichment of genomic variants in PrismNet-predicted high attention regions (HARs) **(A)** Distribution of PhastCons100way conservation scores of HARs, with those of all binding sites from CLIP-seq experiments and all transcript regions as positive and negative controls. **(B)** Enrichment of common SNPs (from 1000 Genomes) in HARs and CLIP binding sites for every RBP comparing to random transcript regions. **(C)** Enrichment of rare SNVs (from 1000 Genomes) relative to common SNVs in HARs and CLIP binding sites for every RBP. All HARs and CLIP-seq binding sites are in K562 cells. In **B** and **C**, each dot represents an RBP. RBPs with HARs significantly depleted/enriched of genomic variants are shown in color (*for P<0.05, permutation or fisher exact test).

**Supplemental Figure 7.**
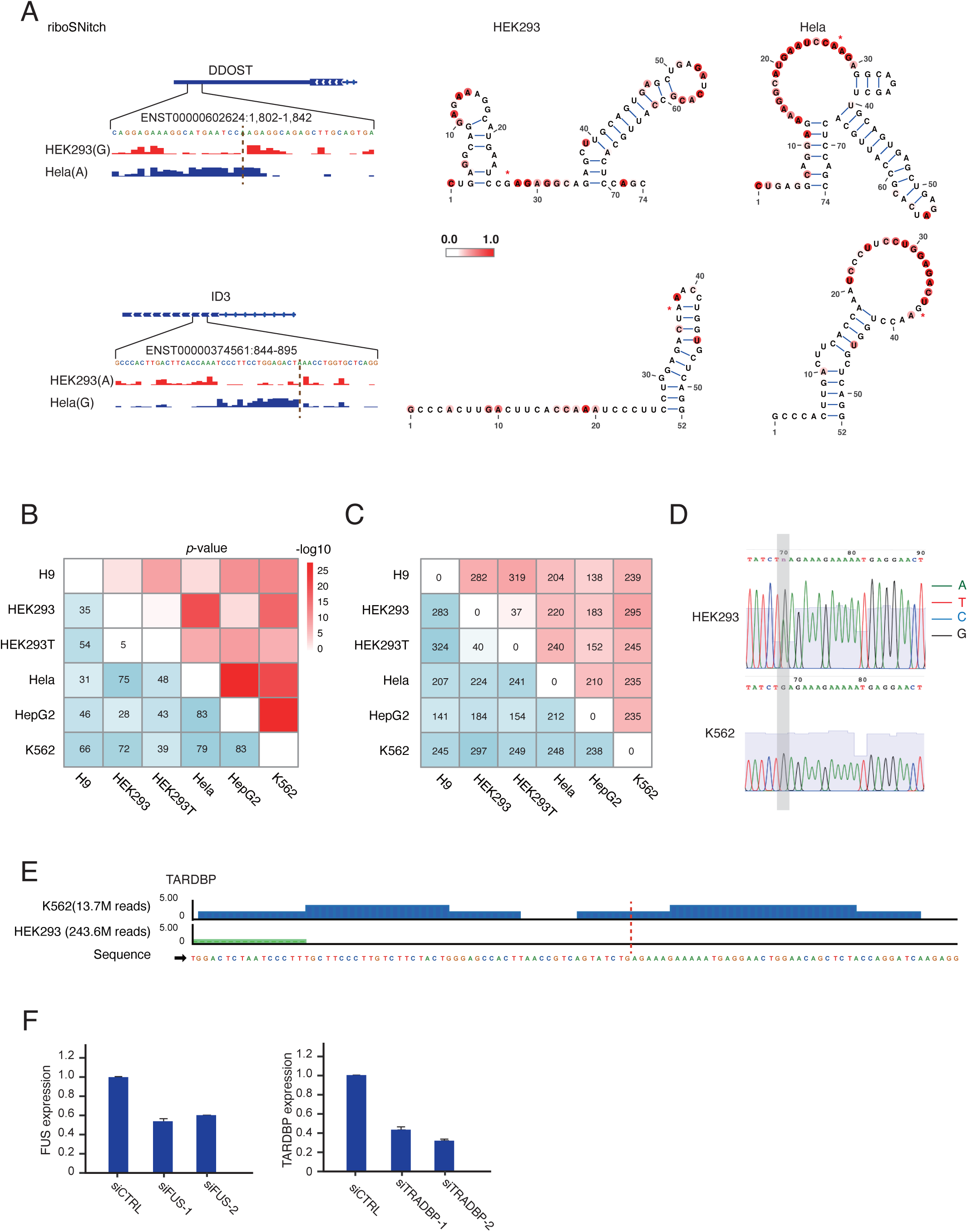
RiboSNitches between different cell lines and their association in RBP dynamic binding sites and human diseases. **(A)** RiboSNitches in the *DDOST* and *ID3* transcripts. Left: Tracks of icSHAPE scores around the riboSNitches in HEK293 and HeLa cell lines. The dash lines indicate the riboSNitch sites. Right: RNA structural models constructed by *RNAshape* with icSHAPE score constraints. **(B)** The number (bottom-left triangle) and the enrichement (up-right triangle) of detected riboSNitches intersecting with the riboSNitches identified in human lymphoblastoid cell lines (Wan et. al., 2014). **(C)** The number of all riboSNitches corresponding to a SNV in ClinVar (bottom-left triangle) and those that are also in dynamic RBP binding sites predicted by PrismNet (up-right triangle). **(D)** Validation of the riboSNitch alleles in the PNPO gene in HEK293 and K562 cells by Sanger sequencing. **(E)** TARDBP binding tracks in K562 (eCLIP) and HEK293 (PARCLIP) in the *PNPO* transcript. The dash line indicates the riboSNitch site. **(F)** The knock down efficiency of FUS and TARDBP in K562 cells.

